# Live 4D monitoring of fungal infection in an ex vivo model of fungal keratitis reveals enhanced efficacy of combined PACK-CXL and natamycin treatment

**DOI:** 10.64898/2026.09.02.748893

**Authors:** Omkaar Sivanesan, Norman Van Rhijn, Shi Ying Tang, Michael J Bromley, Can Zhao

## Abstract

Fungal keratitis (FK) is a major cause of visual impairment worldwide, yet the development and evaluation of new therapeutic approaches are limited by experimental models that inadequately reproduce deep stromal infection or permit longitudinal quantification of fungal burden. Here, we developed an *ex vivo* porcine corneal model of *Aspergillus fumigatus* keratitis that supports reproducible fungal invasion into the deeper corneal stroma and enables non-destructive monitoring of infection over time. Optimisation of the culture conditions demonstrated that reducing nutrient availability markedly promoted stromal penetration compared with nutrient-rich conditions. Using a YFP-expressing *A. fumigatus* strain combined with confocal microscopy and three-dimensional image analysis, fungal burden could be quantified repeatedly within individual corneas, allowing changes in established infection to be monitored longitudinally. We applied the model to evaluate natamycin and photoactivated chromophore for keratitis–corneal cross-linking (PACK-CXL), alone and in combination. Both monotherapies significantly restricted the increase in fungal burden observed in untreated corneas, but neither consistently reduced pre-existing fungal burden. In contrast, combined natamycin and PACK-CXL treatment reduced fungal burden in all treated corneas and was significantly more effective than either treatment alone. Together, these findings establish a quantitative *ex vivo* platform for investigating invasive A. fumigatus corneal infection and demonstrate its ability to distinguish between treatments that restrict fungal expansion and those that reduce established fungal burden. The model provides a useful approach for the longitudinal evaluation and optimisation of therapeutic interventions for fungal keratitis.

## Introduction

Fungal keratitis (FK) is an infection of corneal tissue that can lead to significant vision loss and blindness (Ghenciu et al., 2024; Burton et al., 2011). It has been estimated that more than 1 million people are affected each year with up to 115,000 requiring enucleation (removal of the eye) (Brown et al., 2021). Infections are far more common in tropical and sub-tropical regions; annual incidence in Asia and Africa are estimated at 33.9 and 13.5 cases per 100,000 people compared to 0.02 cases per 100,000 people in Europe (Benedict et al., 2024; Brown et al., 2021; Ong et al., 2016; Limmathurotsakul et al., 2016). Regional aetiology of disease in also differs drastically. Filamentous fungi (*Aspergillus* and *Fusarium* species) are the most common causative fungal organisms in tropical climates (Storr et al., 2024). Comparatively, yeasts (*Candida* species) are more frequently seen in temperate climates (Awad et al., 2024; Ling et al., 2024; Nowik et al., 2020; Tanure et al., 2000).

Topical administration of antifungal medications and oral systemic antifungals are often used as first-line treatment in fungal keratitis management, although therapeutic keratoplasty can also be used in cases with persistent infections (Ansari, Miller and Galor, 2013). The number of FDA approved drugs for fungal keratitis is extremely limited, with natamycin approved for topical treatment only in the USA. Off-label drugs, including the azoles and topical amphotericin B have been used successfully to treat fungal keratitis. However, none of the treatments were found to be superior over others. The antifungal’s ability to infiltrate the corneal tissue presents a limitation of such type of therapy, especially for the treatment of infections at a very late stage and deep within the cornea (Eleiwa et al., 2024; Raj et al.,2021). Fungal keratitis has higher ocular morbidity and mortality outcomes compared to bacterial keratitis, explained by difficulty in diagnosis and reduced effectiveness of available antifungal treatments. Treatment outcomes vary between different regions. In the UK, 44% of the cases were successfully treated with medical intervention (Ting et al., 2021). In the UK and Germany surgical treatment such as penetrating keratoplasty was still needed in 57% and 66% of the cases (Roth et al., 2019; Ting et al., 2021). In the UK, 20% of eyes were blind at final follow up (Ong et al., 2016).

There is a clear and urgent need for alternative effective treatments for fungal keratitis. Previously, researchers have been trying to find innovative therapies beside of antifungal usage for the management of fungal keratitis. Among which, photoactivated corneal crosslinking (PACK-CXL) has been reported as an alternative. PACK-CXL is well established and safely utilised in treatment of corneal infectious conditions caused by bacteria and fungi(Hafezi et al., 2025; Olshaker et al., 2023; Hafezi et al., 2022; Achiron et al., 2021; Sun et al., 2014; Alshehri et al., 2016; Zhu et al., 2018)., with a mechanism by preventing multiplication by intercalating riboflavin, a photosensitizer, into their nucleic acid (Naseem, Ahmad and Hadi, 1988), damaging cell walls by releasing reactive oxygen species (Kumar *et al*., 2004) and preventing secreted enzymes cleavage by alteration of stromal collagen fibril structure (Spoerl, Wollensak and Seiler, 2004). Studies suggested using PACK-CXL treatment alone exhibited successful eradication in wide array of bacterial infections including drug resistant strains, however the results in those cases with fungal originated infections were inconclusive and sometimes even contradictive. The discrepancy between pathogens may be down to fungi more readily invading to deeper corneal layers inaccessible by commonly used treatments. In order to improve the shortcomings of both treatments, combining the PACK-CXL with the topical treatment has recently gained more attention.

A range of experimental systems have been developed to investigate fungal keratitis, including in vitro cell-based approaches and *ex vivo* models using explanted corneal tissue. While *in vitro* systems provide experimentally tractable platforms for investigating specific aspects of host–pathogen interactions, *ex vivo* corneal models retain the three-dimensional tissue architecture required to examine fungal invasion through the corneal stroma. Human corneal tissue has been used for this purpose; however, its limited availability has prompted the development of models using corneas from pigs, rabbits, goats, horses and mice. However, as this resource is exceptionally limited other models using corneas from pigs, rabbits, goats, horses and mice have been developed (Blanco et al., 2017; Hua et al., 2010; Ledbetter et al., 2011; Madhu et al., 2018; Pinnock et al., 2017; Zhou et al., 2011). Irrespective of the *ex vivo* tissue used, previously described fungal infection models have predominantly examined superficial infection, with penetration of fungal rarely extending beyond 100 µm (Alshehri *et al*., 2016). In addition, most of the models described to date require destruction of corneal samples by homogenisation before quantification of infection (e.g. through microbiological culture, qPCR or histology). Thus, these models only allow for single time- point quantification as a proxy to progression of infection. Moreover, histological sections and qPCR cannot reliably differentiate between viable and nonviable fungal cells (Khalid, 2019).

In this study, we have adapted the *ex vivo* model described by Alshehri *et al* to monitor infection with common FK causative agent *A. fumigatus.* We show that by reducing the nutrient content of the aqueous layer used to bathe tissue, we can promote penetration of *A. fumigatus* hyphae into the corneal tissue, enabling the establishment of infections extending into the deeper stroma. By coupling this model with confocal microscopy, we can quantitatively monitor the progression of infection through the corneal tissue. As the isolate employed in this study expresses a cytosolic yellow fluorescent protein (YFP-*A. fumigatus*), fungal structures can be visualised non-destructively within the same cornea at multiple time points. Previous studies have demonstrated that fluorescent protein signal can provide an indicator of fungal viability, with loss of cellular viability associated with a reduction in fluorescent signal (Slade et al., 2013; Young et al., 2010; Webb et al., 2001). YFP fluorescence was therefore used in this study as a proxy for viable fungal burden, enabling longitudinal quantification of changes in fungal burden within individual corneas rather than relying solely on independent endpoint measurements. We subsequently applied this approach to assess changes in fungal burden following treatment with natamycin and PACK-CXL, individually and in combination. Together, this model provides a quantitative ex vivo platform for monitoring the progression of established *A. fumigatus* corneal infection and evaluating treatment-associated changes in fungal burden over time.

## Methods

### Fungal Strains and Media

*A. fumigatus* ATCC46645 expressing cytosolic YFP (PgpdA::yfp(ptrA)) (Lother et al., 2014) was a gift from Prof. Sven Krappmann, and used throughout this study. *A. fumigatus* was cultured on Sabouraud Dextrose agar (Oxoid) for 5 days at 37 °C. Conidia were harvested in phosphate buffered saline + 0.01% Tween-20 (PBS-T) and collected by filtration through Miracloth (Millipore Limited cat. no. 475855). Spores were quantified using a Fuchs-Rosenthal haemocytometer. RPMI-1640 (with L-glutamine, 10% fetal bovine serum (Hyclone, GE Healthcare Life Sciences) and 10% penicillin-streptomycin (PS) (BioWhittaker, USA) was used to culture and maintain the porcine cornea in preparation for fungal infection. Reagents used were sterilised by autoclave or filtration prior to use.

### Porcine Cornea Preparation

Freshly harvested porcine eyes (whole globes) were collected from a local abattoir within 24 hours of porcine sacrifice. Eyes were inspected and any whole globes with evidence of corneal opacity or significant central corneal abrasions were rejected. All subsequent procedures were performed under aseptic conditions. The whole globes were washed with 50 mL of sterile phosphate-buffered saline supplemented with 0.1% Tween-20 and 5% penicillin-streptomycin (PBST-PS, Sigma-Aldrich) for 10 minutes. Corneal dissection was carried out in a Class II microbiological safety cabinet to avoid potential contamination. Using sterile instruments, the cornea was excised from the globe using corneo-scleral dissection at the limbus, leaving a 2-3 mm scleral rim. All other ocular tissue, such as the iris and lens, were removed and disposed. Subsequently, the remaining corneal button was washed 3 times in 5 mL sterile PBST-PS, submerging and moving the button back and forth through the buffer solution for 1 minute. Sterile tweezers were used to hold the scleral rim and taking care not to scratch the corneal surface during this procedure. Washed corneas were then placed in sterile petri-dishes containing fresh PBST-PS as a temporary holding measure until remaining corneas were dissected. Within 1 hour all corneal buttons were dissected and transferred (with the epithelium facing up) into an individual well of a 6-well tissue culture plate containing 1 mL of culture medium (either RPMI-PS or PBST-PS).

### Infection of Cornea with *Aspergillus fumigatus*

The central corneal epithelium of each corneal button was lightly scored using a sterile hypodermic needle (gauge 27G, B Braun™, Fisher Scientific) producing 5 parallel linear abrasions. 1 µL sterile inoculation loops were used to transfer c. 1 µL of inoculum with concentrations between 10^5^ to 10^3^ spore/mL onto corneal surface. The viability of the spores was calculated by performing a viable cell count as described before (Rizzetto et al., 2013) . The 6-well culture plate was covered with lid and left static for 10-15 minutes at room temperature to aid spore attachment to the cornea. Following this, corneal buttons were incubated for the desired period of time (from 1 hour to 48 hours dependent upon experimental setup) at 37 °C, with 5% CO_2_. Uninfected corneas, used as the vehicle control of the experiment, were scored and inoculated with 1 µL of sterile PBS in the same manner, and then maintained alongside the infected corneas in the same 6-well culture plate for each experiment. The culture medium (either RPMI-PS or PBST-PS) was refreshed every 24 hours during the culturing period by first removing the existing medium by pipetting and adding fresh media, avoiding pipette contact with corneal tissue. Gross non-microscopic inspection (viewing the infected well with medium and cornea approximately 15cm away), samples were inspected daily for loss of transparency, discoloration or fur like appearance suggestive of gross fungal growth in the medium to ensure that the only infection site was on the top side of the corneal tissue. Corneal samples with these signs were excluded from further analysis. During each imaging session, corneas were stored at 4 °C within the 6-well plate sealed with additional Parafilm (Bemis, Heathrow Scientific) wrap for <2 hours immediately prior to their imaging window.

### Confocal Microscopy

Cultured corneal buttons were removed using sterile tweezers from the 6-well culture plate and placed with epithelium facing down into 2-well ibidi imaging chambers (ibidi GmbH, Martinsried, Germany). The imaging chamber was filled with 1 mL of medium matching that used for the infection experiments (either RPMI-PS or PBST-PS).

Live cell imaging of the corneal tissue was performed using a Leica TCS SP8 confocal microscope (Leica Microsystems Ltd., Milton Keynes, UK) with a long working distance 25x water immersion objective lens. Excitation and emission wavelength of 514 nm and 525-545 nm respectively were used for imaging the YFP expressed by *A. fumigatus*; the ‘Z-compensation function’ was used for fine adjustment of the laser excitation to compensate the signal reduction in the deeper section of the cornea. These adjustments were done across 5 points of the z-stack, preventing over or under-exposure of acquisition. Acquired images were analysed using Imaris v8.0 software (Bitplane Scientific software module; Zurich, Switzerland). For each cornea, fungal burden was quantified from three independent Z-stacks (466 × 466 µm), each extending to a depth of 1,000 µm through the corneal tissue.

To validate infection depth viewed on microscopy, corneal histology was used as reference comparison. 250ul of 0.1% Cell Mask Deep Red Membrane Stain (Invitrogen™, cat. no. C10046) was used 48 hours post inoculation and allowed to stain cells for 15 minutes prior to microscopic imaging. This marked the corneal epithelium, from which the depth of infection could be accurately measured by disregarding any superficial hyphae growing out from epithelium. From this point, the distance to the deepest fluorescing hyphae was measured producing depth measurement. Microscopically obtained fungal infection depth was compared to the histological depth. Apart from the infection depth validation assay, Cell Mask was not used in other experiments due to its toxic effect.

### Histology

At 36 hours post-infection, each corneal tissue sample was transferred to a fresh well of a 6- well plate containing 2mL of 4% (w/v) paraformaldehyde in PBS (pH 7.4) and kept at room temperature overnight, prior to storage in 70% ethanol. The samples were embedded in paraffin using an automated tissue processor (Leica ASP300, Leica Microsystems). The paraffin wax embedded corneal tissue was then sectioned at 5 *µ*m thickness, using a paraffin microtome (Leica RM2255, Leica Microsystems), onto Superfrost® Plus glass slides (VWR). The sections were dried at 37 °C and then stained with Haematoxylin and Eosin (H&E) or periodic acid Schiff. The slides were left to dry overnight and subsequently stored at room temperature. Images of the stained sections were captured using a slide scanner (3D Histec Pannoramic250 slide scanner, Histec) and analysed using CaseViewer (Histec) software.

### *In vitro* measurement of natamycin’s activity against *A. fumigatus*

Minimum inhibition concentration (MIC) measurement of natamycin was performed as described previously with minor changes (Mendive-Tapia *et al*., 2017). Natamycin were incubated at different concentrations with *A. fumigatus* conidia in a 96-well plate to reach a final volume of 100 µL per well. The final conidia concentration was 5 × 10^5^ cells/mL in RPMI medium. After 48 h incubation at 37 °C, MIC was determined by brightfield microscopy. MIC of natamycin were tested against both *A. fumigatus* strains (MFIG001 and YFP-*A. fumigatus*) (Bertuzzi et al., 2020).

### Comparing the efficacy of the PACK-CXL/natamycin dual therapy with each treatment alone

The *ex vivo* porcine corneal model with *A. fumigatus* keratitis was prepared as previously described. Porcine corneas were infected with ∼1 spore/cornea YFP-*A*. *fumigatus* and the incubated for 36 hours in 1 mL of PBST with 5% penicillin-streptomycin (PBST-PS). Confocal microscopy was used to confirm the establishment of infection. 23 successfully infected corneas were randomly allocated into 4 groups: control group with PBST treatment (5), PACK- CXL treatment group (6), natamycin treatment group (6) and PACK-CXL/natamycin dual treatment group (6). All samples were kept under the same conditions (room temperature, inside of a Class II cabinet) during the treatment period, and were then transferred into a fridge to be kept at 4 °C while the confocal microscopy being conducted.

PACK-CXL treatment was carried out as described per Dresden protocol (Wollensak, Spoerl and Seiler, 2003) in a Class II cabinet to avoid potential contamination. The corneal button was mounted onto an artificial chamber with retainer (Figure 1). PBST-PS was infused to fill the space underneath the cornea using a 15 mL syringe via the tubing system connected to the chamber, mimicking the *in vivo* intraocular environment. A sterile blunt scalpel was used to debride the corneal epithelium, followed by irrigation using ddH_2_O. With the light switched off, isotonic riboflavin eye drops (0.1% [wt/wt] riboflavin, 20% [wt/wt] dextran T500) was topically administrated every 5 minutes within a 30-minute time frame using a pipette to cover the entire corneal surface. The prepared to cornea was then irradiated for 30 minutes with an UV-A radiation of 5.4J/cm^2^ using a VEGA clinical cross-linking machine (370 nm, irradiance of 3 mW/cm^2^, Florence, Italy), with intermittent riboflavin administration again at 5-minute intervals. After treatment, the cornea was dismounted, rinsed with PBS and then incubated in a lidded 6-well plate with 1 mL fresh PBST at room temperature for a further 6- hour period, while the Natamycin treatment being conducted on the other experimental groups.

**Figure 1.**
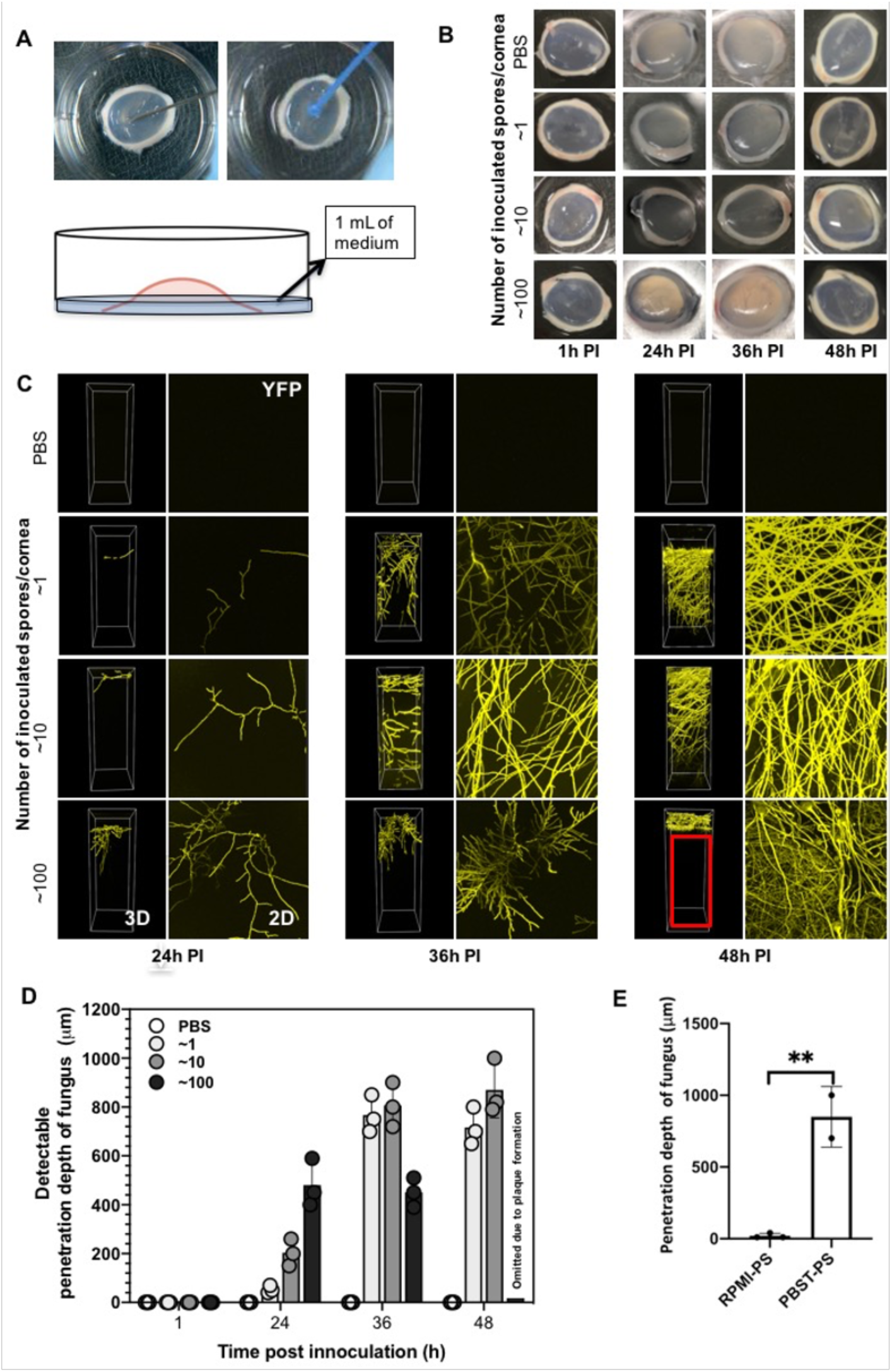
*A. fumigatus* infections developed in corneas inoculated with 4 different inoculum levels incubated at 37 °C for different durations in PBST-PS. (A) A schematic diagram of the ex *vivo* infection model. (B) Images of corneas at 48 hours post inoculation *of A. fumigatus* spores at different levels (∼1, ∼10 and ∼100 spores/cornea) and PBS after different incubation periods. (C) Live cell imaging through the corneas with epithelium surface at the top in 3D projections (left panel), and superimposed birds eye view through the cornea (right panel), yellow signal represent live fungal filaments of cytoplasm YFP expressing *A. fumigatus.* Red rectangular indicates where fluorescent signal was blocked by plaque. (D) Mean penetration depth values were determined using Imaris v8.0 and represented as means±SD (n=3). The effect of time and inoculum on penetration depth was assessed by Two-way ANOVA. (E) Corneas were infected with ∼1 spore/cornea and then incubated for 48 hours in either RPMI-PS or PBST-PS. Mean penetration depth values were determined using Imaris v8.0 and represented as means +SD. Statistical difference was assessed by student T-test (** equals p< 0.01, n = 3 cornea)

Natamycin treatment was carried out in a Class II cabinet at room temperature. Corneas were placed in a fresh 6-well plate along with 1 mL fresh PBST-PS, leaving the corneal surface exposed in the air while the remaining part submerged in PBST. 20 µL of natamycin solution was topically administrated onto the corneal surface at 2-hour intervals for a 6-hour period (4 treatments in total). Natamycin was used at 120 mg/L, which was the highest soluble concentration can be used without any precipitation during the experimental period. Corneas were kept completely stationary during the whole treatment period, to avoid the natamycin droplet from falling off from the central corneal area into the PBST-PS. After the treatment, corneas were rinsed with PBS to remove the excess of natamycin, and were then transferred into a fresh 6-well plate with PBST-PS and kept in a fridge at 4 °C for a short period while the confocal microscopy being conducted.

For the PACK-CXL/natamycin dual treatment, PACK-CXL treated corneas were treated with natamycin solution immediately as described above. After the treatment, corneas were rinsed and kept in a 6-well plate at 4 °C as the other experimental groups while the confocal microscopy being conducted.

Confocal microscopy was performed immediately before treatment and again following the treatment period in all experimental groups, with consistent acquisition and analysis settings applied across all time points. Fungal burden was quantified for each individual cornea before and after treatment, allowing the change in fungal burden within the same cornea to be determined.

### Data analysis and Quantification

Fluorescence-based quantification was performed using two complementary approaches depending on the experimental stage. During initial development and optimisation of the infection model, YFP fluorescence was quantified using an area-under-the-curve (AUC)-based approach. Unless otherwise stated, RenyiEntropy was used for automatic thresholding (Kapur, Sahoo and Wong, 1985). Thresholded Z-stacks were quantified using a macro, whereby individual images within each stack were converted to binary images and the number of YFP-positive pixels within each optical section was determined. YFP-positive signal was plotted as a function of imaging depth using ggplot2 (Wickham, 2016), and the AUC of the resulting fluorescence-depth profile was calculated as a measure of fungal burden. This approach was used during model optimisation to compare the extent and distribution of fungal infection between experimental conditions and time points.

For subsequent treatment experiments, fungal burden was quantified by three-dimensional reconstruction of confocal Z-stacks using Imaris v8.0 (Bitplane Scientific Software, Zurich, Switzerland). YFP-positive fungal structures were segmented using the ‘Surface’ module, and the volume occupied by the fluorescent signal was calculated to provide a volumetric measure of fungal burden. Consistent segmentation parameters were applied across all samples within each experiment. As individual corneas were imaged immediately before and following treatment, treatment response was determined from the change in fungal burden within each cornea between the paired time points.

## RESULTS

### A novel *ex vivo* porcine model of FK supports deep stromal invasion and longitudinal monitoring of A. fumigatus infection

To enable adequate evaluation of novel antifungal treatments, experimental models of FK should reproduce the clinically relevant feature of fungal invasion into the deeper corneal stroma. We therefore assessed the depth of infection in porcine corneal buttons that had been inoculated with *circa* 100 spores of an *A. fumigatus* isolate that expresses a cytoplasmic yellow fluorescent protein (YFP) following a method previously developed for *Fusarium oxysporum* infections (Alshehri *et al*., 2016). Forty-eight hours post infection the corneal surface was covered by a visible opaque plaque consisting of a matt of hyphae and pigmented conidiophores (Supplemental Figure 1A). Attempts were made to assess fungal penetration into the cornea using live-cell fluorescent confocal microscopy (Supplemental Figure 1B). However, the extensive fungal biomass on the corneal surface prevented light penetration to deep corneal layers, masking the depth of infection (Supplemental Figure 1C and D)

With the aim of improving our ability to evaluate fungal penetration in the cornea by reducing growth on the surface, the inoculum level was reduced to *c.*10 and *c.*1 spore per cornea. After 48 hours, all inoculated corneas (n=3 at c. 10 spores per inoculum; n=3 at c. 1 spore per inoculum) developed *A. fumigatus* infections, while control cornea (PBST) remained clear of infection. Corneas infected with ∼1 spore developed an infection that could not be seen macroscopically (Supplemental Figure 1A). Fluorescent confocal microscopy revealed the presence of an extensive network of fungal filaments on the surface of the cornea, but the infection was superficial, not penetrating beyond 30 *µ*m of the corneal surface (Supplemental Figure 1B).

In nutrient-rich culture media such as RPMI-1640, *A. fumigatus* downregulates pathways involved in carbon and nitrogen acquisition, including proteases that may contribute to tissue invasion (Farrell *et al*., 2017; Ries et al., 2019). We therefore tested whether the limited stromal penetration observed under RPMI-1640 culture conditions could be improved by reducing nutrient availability in the aqueous phase (Figure 1A). Replacing RPMI-1640 with PBS containing 0.01% Tween-20 (PBST) resulted in a marked increase in fungal penetration depth after 48 h, from 7 ± 3 µm in RPMI-1640 to 783 ± 214 µm in PBST (P < 0.01; Figure 1E). These data indicate that nutrient limitation within the culture system promotes deeper stromal invasion by *A. fumigatus*.

Having established PBST as the preferred culture condition for promoting stromal invasion, we next examined the relationship between inoculum size and infection duration. 12 corneas were inoculated with approximately 1, 10 or 100 conidia and infection progression was monitored over time. Despite the low inoculum, all infected corneas developed detectable infection by 24 h, while the vehicle control remained negative (Figure 1C). Penetration increased over the first 36 h, with the deepest invasion observed in corneas inoculated with approximately 1 or 10 conidia (Figure 1D). By 48 h, infections initiated with 1 or 10 conidia continued to show increased fungal burden, while penetration depth remained broadly similar to that measured at 36 h. In contrast, the 100-conidium condition produced a dense surface plaque by 48 h, resulting in an artefactually low depth measurement because of reduced optical penetration (Figure 1C,D). Taken together, these observations indicated that an inoculum of approximately 1–10 conidia per cornea combined with an incubation period of 36–48 h provided the most suitable conditions for establishing deep stromal infection while retaining the ability to monitor infection by confocal microscopy.

When opting for ∼1 spore per cornea there is a risk of not transferring any inocula onto the cornea. However, a lower initial infection burden allows similar areas within the same cornea to be repeatedly identified and assessed to monitor progression of infection closely and mimics real-world infections scenario. Using a lower inoculi also enable us to monitor the cornea longer than with higher inoculi. In order to mitigate such risks when using a low inoculi (∼1 spore per cornea), we assessed the reproducibility of our model and also the analytical pipeline. We replicated the infection model across nine independent experiments, with 10 corneas included in each experiment (90 corneas in total), using the established conditions of 36 hours post infection in PBST. After 36 hours, 66 corneas out of 90 developed *A. fumigatus* infection giving an average successful infection rate of 73% (adjusted Wald confidence intervals 63-81%) (Figure 2C; Supplemental Table 1). The overall average depth of infection of 559 ±26 *µ*m with no significant differences between the nine independent experiments (Supplemental Figure 3). Of the successful infections, 83.3% (55 out of 66) developed an invasive infection that penetrated greater than 400 *µ*m from the corneal surface (Figure 2D). Measurement of corneal fungal burden from 9 representative corneas between 24 hours and 36 hours post infection, quantified as average area under the curve (AUC) of the fluorescent signal, revealed a time-dependent increase of fungal burden across the corneal tissue over the 12 hours period (Figure 2A and B) (p < 0.01). These results demonstrated that, despite the low starting inoculum, the selected conditions were capable of consistently generating deep stromal infection in corneas in which infection was successfully established.

**Figure 2.**
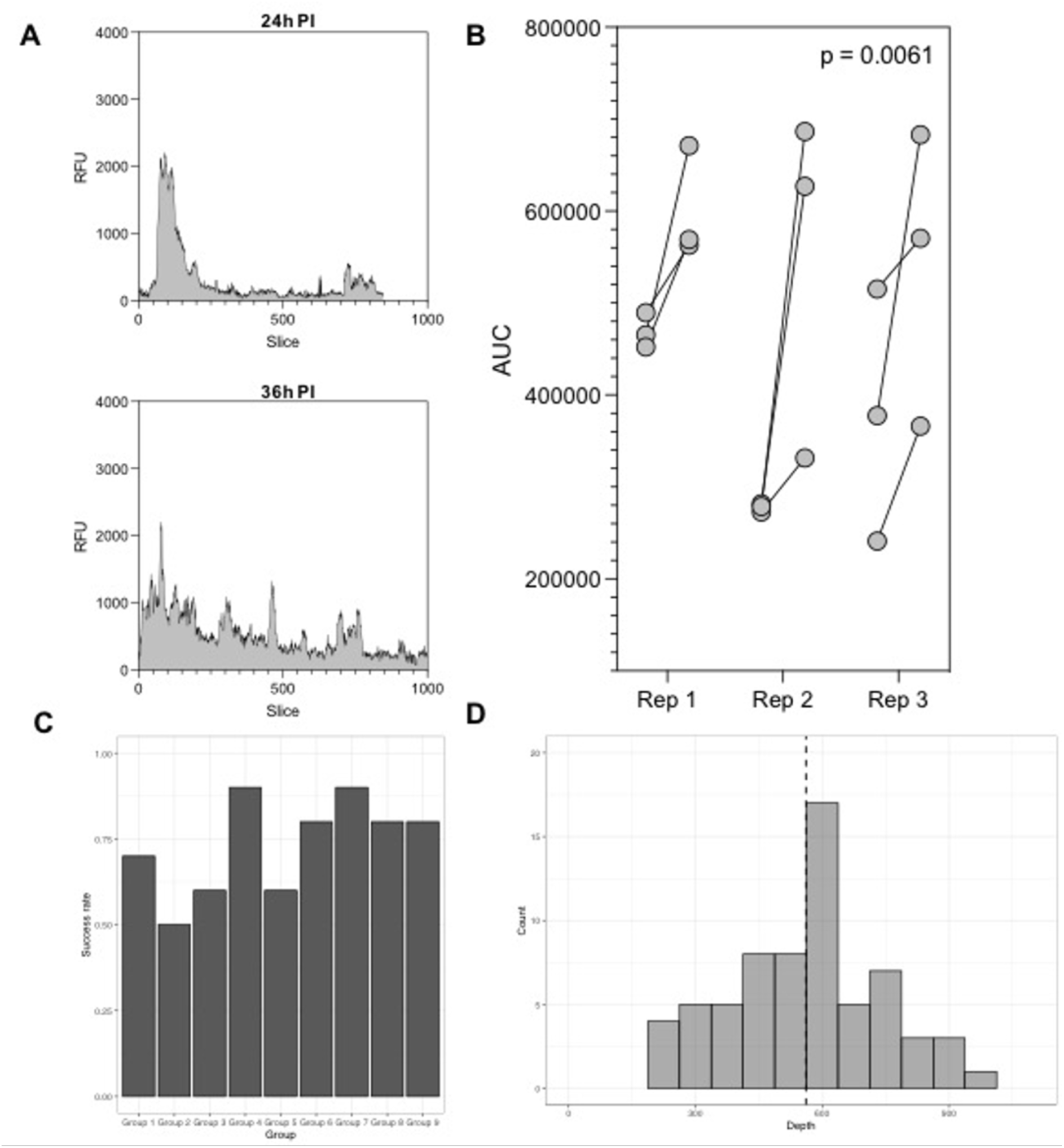
Quantification of fungal burden within the corneal tissue and assessment of the reproducibility of the infection model. (A) Relative fluorescent units (RFU) determined from a representative cornea infected with 1 spore at 24 hours and 36 hours post infection. (B) Area under the curve (AUC) quantification of 9 corneas infected with 1 spores at 24 hours and 36 hours post infection. (C) Success rate of establishing a deep seated infection for each of the experimental group (n=9 per group). (D) penetration depth of the fungus 36 hours post-infection as measured by the maximum depth of which fluorescent intensity was higher than background signal. The mean depth of penetration was 558.9 pm.

To independently confirm that the YFP-positive fungal structures detected by confocal microscopy represented stromal invasion, infection depth was additionally assessed by histological examination. Fungal hyphae were observed extending through the corneal stroma, with the deepest hyphae detected at approximately 1754–1952 *µ*m in the representative samples examined. These observations confirmed that *A. fumigatus* was capable of penetrating extensively through the corneal tissue under the optimised culture conditions and supported the use of fluorescence confocal microscopy for monitoring stromal invasion in the intact cornea.

### *Ex vivo* porcine corneal infection model can be used to assess antifungal treatment efficacy in real-time

To assess the feasibility of the ex vivo corneal model for evaluating antifungal treatment efficacy, corneas were infected with *A. fumigatus* and incubated for 36 h to allow stromal infection to become established. Successful infection was confirmed by confocal microscopy before treatment (Figure 3A, before treatment), after which corneas were treated with PBST (Untreated), natamycin, PACK-CXL, or a combination of natamycin and PACK-CXL. The same corneas were subsequently re-imaged 6 h after treatment commenced (Figure 3A, after treatment). Three-dimensional reconstruction and volumetric analysis of YFP-positive fungal structures allowed fungal burden to be quantified before and after treatment within each individual cornea, enabling treatment-associated changes in fungal burden to be assessed longitudinally.

**Figure 3.**
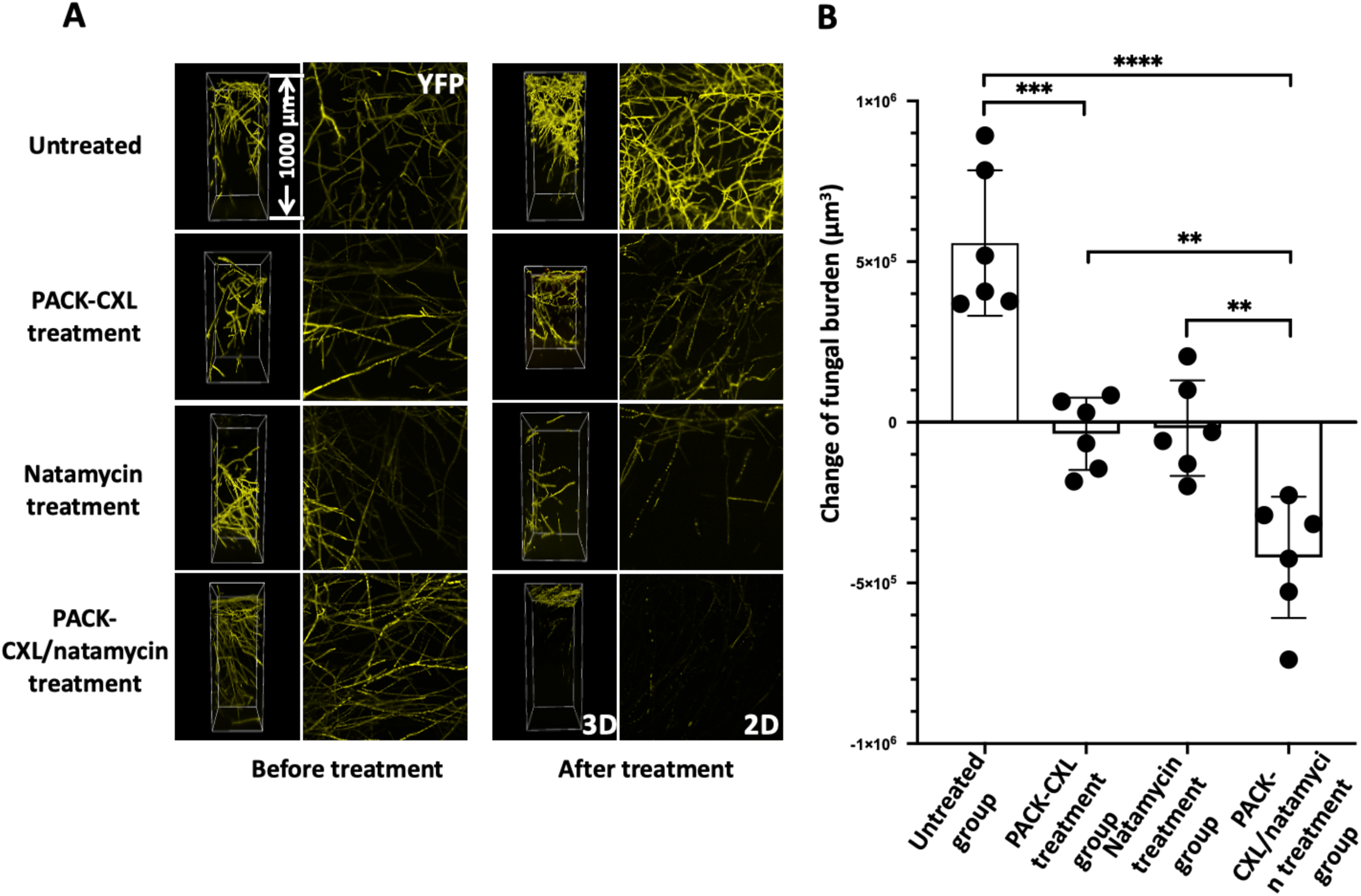
Evaluation of the efficacy of PACK-CXL/natamycin dual treatment compared to each treatment alone. (A)Live cell imaging through the corneas with epithelium surface at the top in 3D projections (left panel), and superimposed birds eye view through the cornea (right panel), yellow signal represent live fungal filaments of cytoplasm YFP expressing A. fumigatus. (B) Scatter dot plot showing mean change in fungal burden with SD for each treatment group and one-way ANOVA comparison of groups indicated by stars of significance. Each dot stands for the mean change of occupational volume (pm3) from 3 scanning within one cornea from the treatment group. The mean change of the fungal burden within each cornea sample were calculated by deducting the initial occupational volume before the treatment from the occupational volume obtained after the treatment. **\* = p < 0.05, ** = p < 0.01, *** = p < 0.001, **** = p < 0.0001.**

Fungal burden increased in all PBST-treated control corneas during the 6 h experimental period, consistent with continued progression of the established infection (Figure 3B). Both PACK-CXL and natamycin significantly restricted this increase compared with the PBST-treated control group (P < 0.0001 for both comparisons). However, responses to either monotherapy were heterogeneous, with both increases and decreases in fungal burden observed between individual corneas, and there was no significant difference between PACK-CXL and natamycin treatment (P = 0.9981). Thus, under the conditions examined, both monotherapies predominantly limited further expansion of the established fungal burden rather than producing a consistent reduction from pre-treatment levels. In contrast, combined treatment with PACK-CXL and natamycin resulted in a reduction in fungal burden in all six treated corneas and was significantly more effective than either PACK-CXL alone (P = 0.0054) or natamycin alone (P = 0.0037), as well as the PBST-treated control (P < 0.0001) (Figure 3B). These findings demonstrate that the model can distinguish between treatments that restrict the progression of established infection and those that produce a measurable reduction in fungal burden over time.

## DISCUSSION

Fungal keratitis is a major cause of visual impairment and blindness globally (Wong et al., 1997), and effective management depends on the timely control and eradication of fungal infection. Experimental models that reproduce key features of corneal infection while allowing fungal progression and treatment response to be quantitatively assessed are therefore important for the development and evaluation of new therapeutic approaches. In this study, we established an *ex vivo* porcine corneal model that supports reproducible *A. fumigatus* invasion into the deeper corneal stroma and permits non-destructive, longitudinal monitoring of infection within individual corneas. By combining a YFP-expressing *A. fumigatus* strain with confocal microscopy and three-dimensional image analysis, changes in fungal burden can be quantified over time within the same infected tissue. Importantly, this approach also enables treatment-associated changes in established fungal burden to be measured, allowing the response to different antifungal interventions to be compared longitudinally.

Previous *ex vivo* models of fungal keratitis have commonly employed nutrient-rich culture media, including Eagle’s medium, to maintain infected corneal tissue (Hua et al., 2010; Ledbetter, Irby and Kim, 2011; Zhou et al., 2011; Alshehri et al., 2016; Blanco et al., 2017; Pinnock et al., 2017; Madhu et al., 2018). In the present study, however, culture under nutrient-rich conditions predominantly supported superficial growth of *A. fumigatus*, with limited penetration into the corneal stroma, and reducing the initial inoculum alone was insufficient to overcome this phenotype. In contrast, replacing RPMI-1640 with the relatively nutrient-limited PBST-PS resulted in a marked increase in stromal penetration, indicating that nutrient availability within the culture system is an important determinant of the infection phenotype generated by this *ex vivo* model. One possible explanation is that readily available nutrients in the surrounding medium favour fungal proliferation at the corneal surface, whereas nutrient limitation may promote fungal invasion into the tissue in search of alternative nutrient sources. However, the mechanisms underlying this response were not investigated directly in the present study. From a practical perspective, the use of PBST-PS therefore provided conditions that more reliably supported the establishment of deep stromal infection, which was required for subsequent longitudinal imaging and treatment studies.

A further practical advantage of the model is that specialised equipment is not required to establish infection. During incubation, corneas are maintained in standard lidded culture plates using an air–liquid interface, with the epithelial surface exposed to air while the endothelial surface remains in contact with the culture medium. This approach is based on the air–liquid corneal culture system originally described by Richard et al. (1991), which was developed to better reproduce the physiological environment of the cornea.

Porcine corneas were selected for this model because they are readily obtainable from animals slaughtered for the food industry and share several anatomical and physiological characteristics with the human cornea. Compared with rodent corneas, porcine corneas more closely resemble human corneas in their three-dimensional architecture and collagen and extracellular matrix composition, and porcine corneal tissue has previously been investigated for xenotransplantation (Hara and Cooper, 2011). Nevertheless, important differences between porcine and human corneas remain. Porcine corneas are generally thicker, with reported thicknesses ranging from 666–1,013 µm, and differ in parameters including diameter and astigmatism (Jay et al., 2008; Sanchez et al., 2011). Bowman’s layer also appears to be absent from the porcine cornea, and differences in corneal water content have been reported (Sanchez et al., 2011; Taylor et al., 2015). These differences should be considered when extrapolating observations from the present model to human corneal infection. Despite these limitations, porcine corneas are widely used in *ex vivo* vision research and provide an accessible tissue system with substantial structural similarity to the human cornea (Sanchez et al., 2011).

Notably, we did not observe corneal ulceration or oedema during infection, even in corneas exhibiting extensive fungal growth, consistent with observations reported in a *previous ex vivo* corneal infection model (Pinnock et al., 2017). This likely reflects an important limitation of the ex vivo system: although the corneal tissue architecture is retained, the model lacks an intact host immune and inflammatory response. In vivo, the development and progression of fungal keratitis involve complex interactions between the invading fungus, resident corneal cells and recruited immune cells. Inflammatory responses contribute substantially to the tissue pathology associated with keratitis, including epithelial damage and ulceration (Ruiz-Ederra et al., 2005; Oguz et al., 2005), stromal polymorphonuclear neutrophil infiltration (Shetty et al., 2014; Hsiao et al., 2015), and ulcer formation (Sueke et al., 2013). Consequently, the present model is principally suited to examining fungal growth and invasion within intact corneal architecture, and treatment-associated changes in fungal burden, rather than reproducing the complete host–pathogen interaction or inflammatory pathology of fungal keratitis.

Under the optimised experimental conditions, fungal infection could be visualised for up to 48 h following inoculation with approximately one conidium per cornea, an inoculum substantially lower than those employed in previously reported ex vivo models. Importantly, the model supported reproducible fungal invasion into the deeper corneal stroma while retaining sufficient optical transparency for longitudinal confocal imaging. The use of YFP-expressing *A. fumigatus* further allowed fungal burden to be quantified non-destructively at multiple time points within the same cornea. Consequently, rather than relying solely on independent endpoint measurements, changes in fungal burden and stromal invasion can be followed within individual infected corneas as infection progresses. This provides a quantitative approach for examining the development of invasive *A. fumigatus* infection within intact corneal tissue and, importantly, for measuring subsequent changes in established fungal burden following therapeutic intervention.

An application of this longitudinal approach is the assessment of treatment response in established infection. Using paired measurements of fungal burden obtained from individual corneas before and after treatment, we were able to distinguish continued infection progression in untreated corneas from the responses observed following natamycin, PACK-CXL, or combined therapy. Both natamycin and PACK-CXL significantly restricted the increase in fungal burden observed in untreated corneas (P < 0.0001 for both comparisons), although neither monotherapy produced a consistent reduction in pre-existing fungal burden across individual corneas. In contrast, combined natamycin and PACK-CXL treatment reduced fungal burden in all six treated corneas and produced a significantly greater effect than either PACK-CXL (P = 0.0054) or natamycin (P = 0.0037) alone. These findings suggest that, under the conditions examined, either monotherapy was sufficient to restrict further expansion of established infection, whereas the combination produced a more consistent reduction in the fungal burden already present at treatment initiation.

Natamycin was used at 120 mg/L, corresponding to 30-fold its MIC (4 mg/L) against the A. fumigatus strain used and representing the highest concentration that could be maintained in solution without visible precipitation under the experimental conditions. This differs from the 5% ophthalmic suspension used clinically and should therefore be considered when interpreting the treatment response observed in this model. The present experiments were designed to assess the ability of the model to quantify treatment-associated changes in fungal burden rather than to reproduce the clinical pharmacokinetics or dosing of natamycin.

The combined treatment with natamycin and PACK-CXL produced the most pronounced response observed in this study. Whereas either treatment alone significantly restricted the progression of fungal burden relative to untreated controls, neither monotherapy consistently reduced the fungal burden present at treatment initiation. In contrast, combined treatment resulted in a reduction in fungal burden in all treated corneas and was significantly more effective than either treatment alone. Importantly, residual fungal burden remained detectable following treatment, and the observed reduction should therefore not be interpreted as complete eradication of infection. Rather, these findings demonstrate that the longitudinal nature of the model can distinguish between inhibition of further fungal expansion and a measurable reduction in established fungal burden. While the absence of an intact immune response and other features of the living ocular environment limits direct extrapolation to clinical treatment, the model provides a reproducible and quantitative platform for investigating invasive *A. fumigatus* infection within corneal tissue and for comparing treatment responses over time. Further studies will be required to determine how observations made using this *ex vivo* system translate to in vivo infection; nevertheless, the ability to repeatedly quantify established infection within the same cornea provides a useful intermediate approach for the evaluation and optimisation of antifungal interventions.

---

**Supplemental Figure 1.**
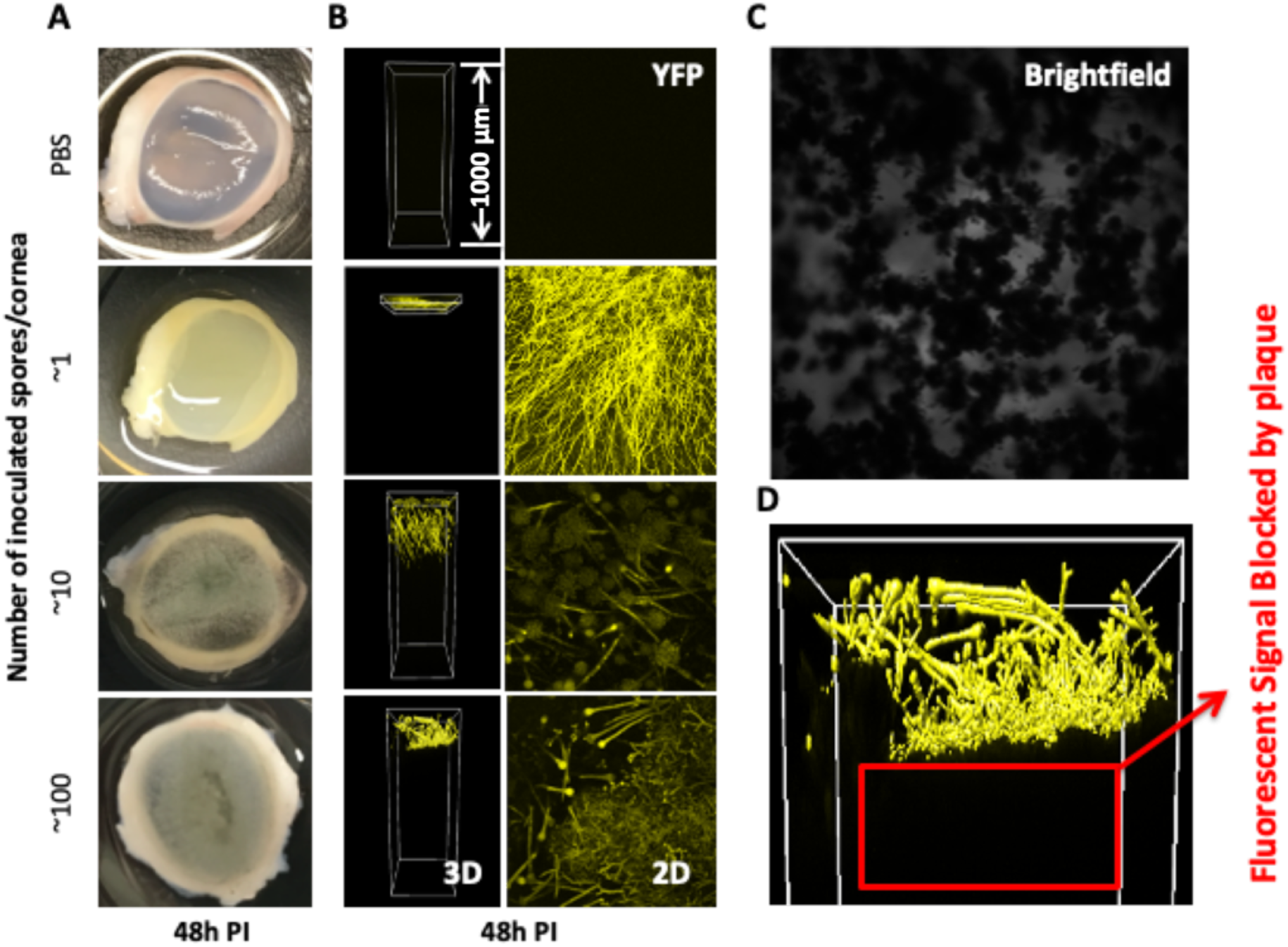
***A. fumigatus* infections developed in corneas inoculated with 4 different inoculum levels after 48 hours incubation at 37 °C in RPMI-PS** (A) Images of corneas at 48 hours post inoculation *of A. fumigatus* spores at different levels (∼1, ∼10 and ∼100 spores/cornea) and PBS. (B) Live cell imaging through the corneas with epithelium surface at the top in 3D projections (left panel), and superimposed birds eye view through the cornea (right panel), yellow signal represent live fungal filaments of cytoplasm YFP expressing *A. fumigatus.* (C) Brightfield image of a plaque formed on cornea infected with ∼100 spores at 48 hours post inoculation in RPMI-PS. (D) Fungal plaque blocking fluorescent signal being detected through the scan from epithelium surface.

**Supplemental Figure 2.**
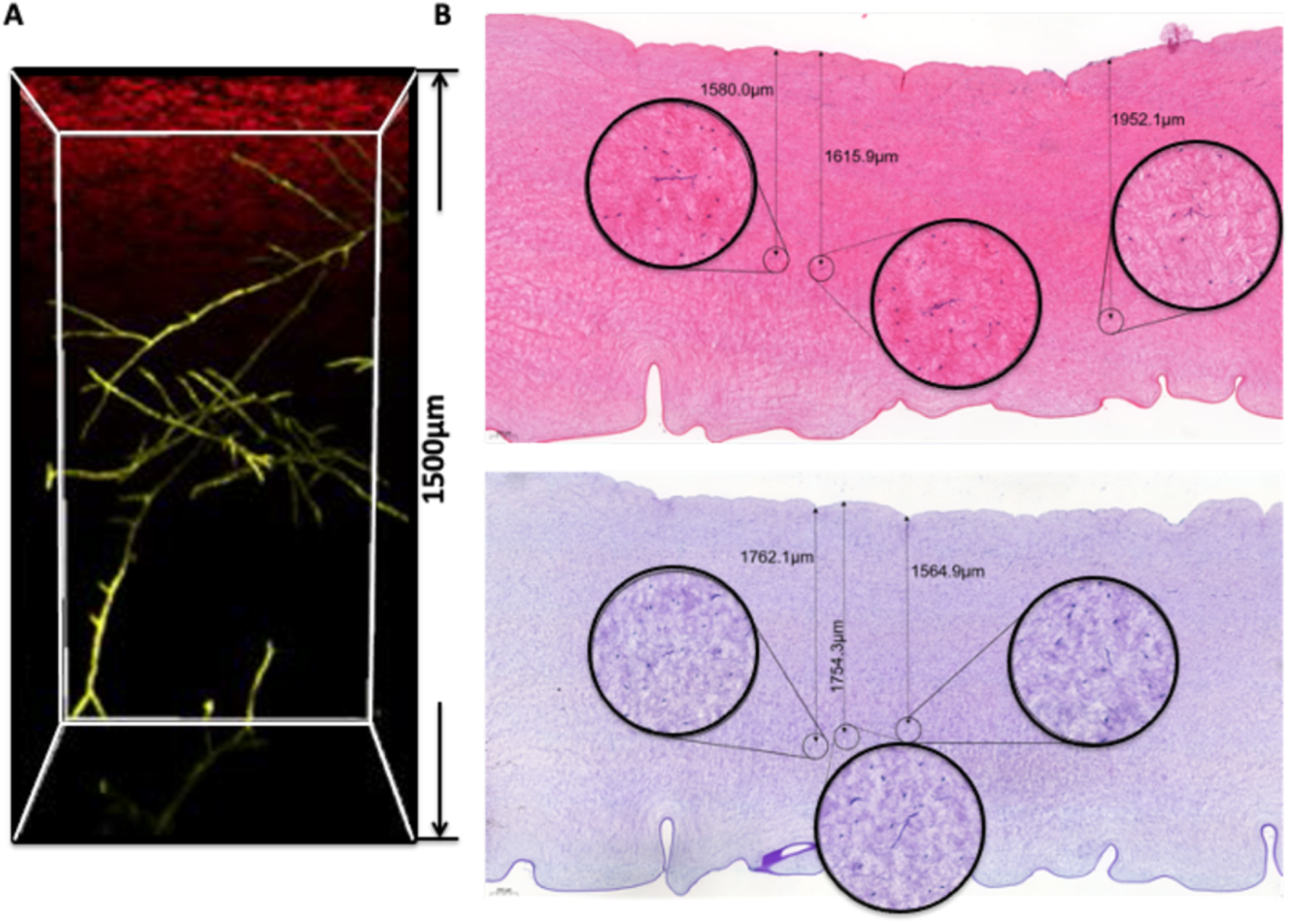
Histology of *A. fumigatus* infection of *ex vivo* porcine cornea confirmed progression of infection visualised using confocal microscopy. (A) Fluorescent confocal images of infected cornea at 48 hours post inoculation of ∼1 *A. fumigatus* spores at different levels. Red fluorescent signal indicate corneal surface stained by 0.1% Cell Mask Deep Red Membrane Stain, and yellow signal represent live fungal filaments of cytoplasm YFP expressing *A. fumigatus.* (B) Corneas were fixed in 10 % buffered formalin, embedded in paraffin, sectioned and stained with haematoxylin and eosin stain. *A. fumigatus* filaments can be observed at the epithelial surface and are present in the stroma. Three deepest penetrated filaments are enlarged, with distance from corneal surface to filament shown.

**Supplemental Figure 3.**
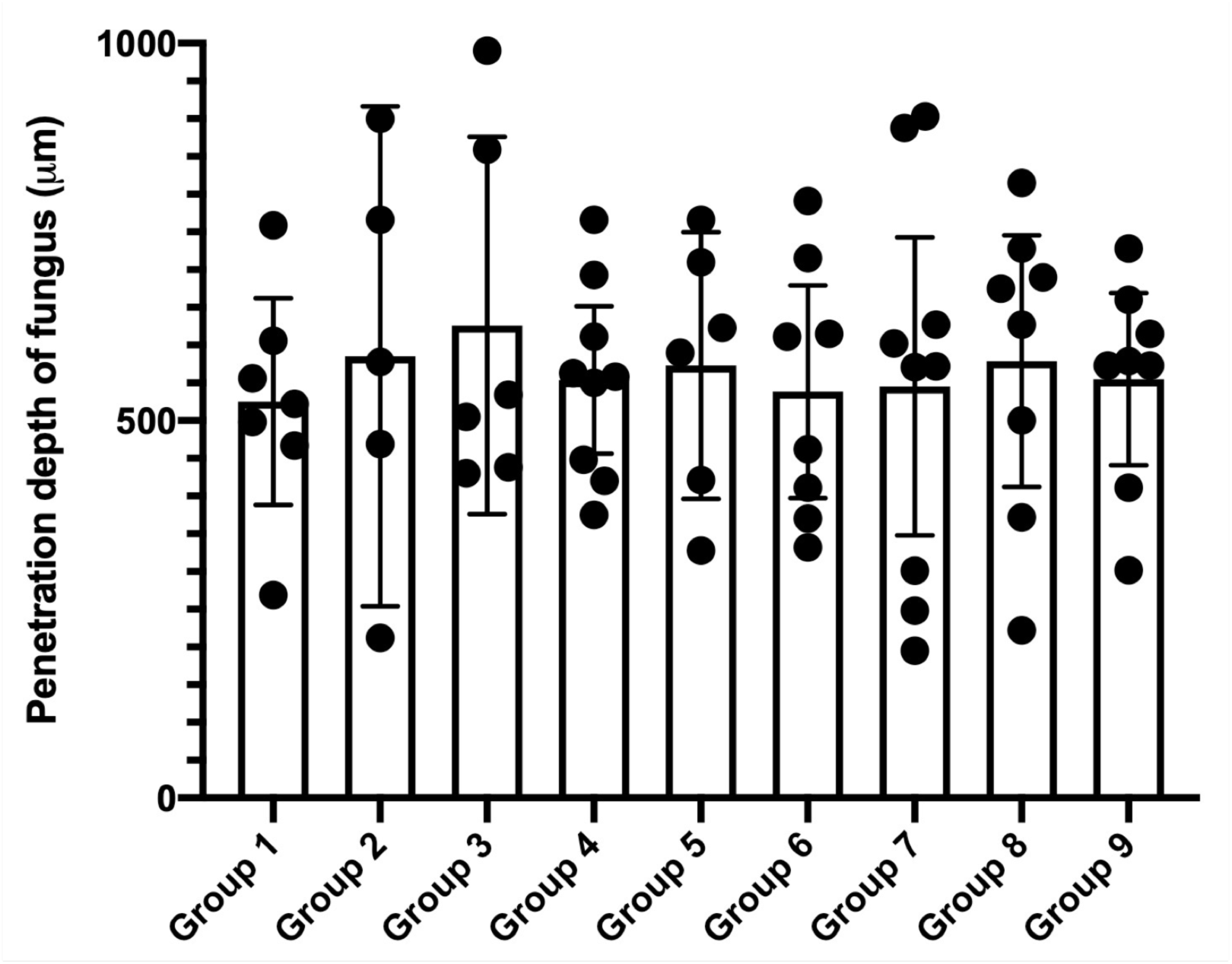
Reproducibility of a low inoculum ex vivo keratitis model. Plot with all corneal replicates successfully infected with ∼1 spore/cornea after 36 hours incubation period in PBST-PS before confocal microscopy., represented by black circles. Uninfected corneas were omitted from the plot. Mean penetration depth values were determined using Imaris v8.0 and represented as means ±SD. Statistical difference was assessed by ANOVA, no statistical difference between groups was found (n = 10 cornea per group).

**Supplemental Table 1.** Reproducibility of ex *vivo* porcine corneal infection after 36 hours incubation with ∼1 spore/cornea inoculum in PBST-PS

|  | Group 1 | Group 2 | Group 3 | Group 4 | Group 5 | Group 6 | Group 7 | Group 8 | Group 9 |
| --- | --- | --- | --- | --- | --- | --- | --- | --- | --- |
| Penetration depth (µm) |  |  |  |  |  |  |  |  |  |
| Cornea 1 | 269 | NI | NI | 766 | 328 | 370 | 888 | 500 | 660 |
| Cornea 2 | 498 | NI | 990 | 420 | 766 | NI | 602 | 690 | 573 |
| Cornea 3 | 467 | 469 | NI | 558 | 710 | 411 | 571 | NI | 615 |
| Cornea 4 | NI | NI | NI | 375 | 623 | 462 | 903 | 815 | 302 |
| Cornea 5 | 556 | 578 | 430 | 448 | NI | 715 | 195 | 675 | 728 |
| Cornea 6 | 606 | 212 | 859 | 611 | NI | 791 | 627 | 728 | NI |
| Cornea 7 | 522 | NI | 438 | 693 | NI | 332 | 301 | 372 | 579 |
| Cornea 8 | 759 | 900 | 534 | 550 | 421 | NI | 248 | NI | 573 |
| Cornea 9 | NI | 766 | NI | NI | NI | 611 | 572 | 627 | 411 |
| Cornea 10 | NI | NI | 505 | 563 | 590 | 615 | NI | 222 | NI |
| Success rate of Infection of each group (%) | 70 | 50 | 60 | 90 | 60 | 80 | 90 | 80 | 80 |
| Average penetration depth of each group (µm) | 525.2 | 539.7 | 650.2 | 552.6 | 569.6 | 513.5 | 545.2 | 578.6 | 555.1 |
| Average Success rate of Infection (%) |  |  |  |  |  | 73.3 |  |  |  |
| Average infection depth with success infection (µm) |  |  |  |  |  | 558.9 |  |  |  |

